# *TaWUS2D* regulates the number of grains per spikelet by enhancing the number of fertile ovaries in Multi-Ovary Wheat

**DOI:** 10.1101/2025.02.05.636547

**Authors:** Adam Schoen, Parva Kumar Sharma, Alex Mahlandt, Andy Chen, Huajin Sheng, Leon Kochian, Peng Gao, Daoquan Xiang, Teagen Quilichini, Prakash Venglat, Sheng Wang, Inderjit Singh Yadav, Yong Gu, Daniel Rodriguez-Leal, Weifeng Luo, Yiping Qi, Nathan Meier, Anmol Kajla, Matthew Willman, Gina Brown-Guedira, Cal Youngblood, Amanda Hulse-Kemp, Angus Murphy, Bikram Gill, Cristobal Uauy, Raju Datla, Nidhi Rawat, Vijay Tiwari

## Abstract

Innovative genetic improvements in the staple crop *Triticum aestivum* (bread wheat) are urgently needed to address the growing global food security crisis. Here, we report the map-based cloning of *TaWUS2D*, the gene responsible for the dominant multi-ovary phenotype in wheat. Multi-ovary lines are characterized by the development of three fertile ovaries per floret that results in three grains, as opposed to wildtype single ovary wheat. We used HiFi long-reads to assemble a 14.48 Gbp genome scaffold assembly in the background of mutli-ovary wheat line MOV. Using high-resolution genetic mapping, combined with additional genomic resources, we defined the *Mov-1* locus to a 135 Kbp region containing two genes. Using five independent deletion mutants and eight TILLING mutants, we demonstrate that a functional WUSCHEL-like protein, *TaWUS2D*, is required for the multi-ovary phenotype. *TaWUS2D* is upregulated in the MOV genetic background. This research lays the groundwork for developing new approaches to improve wheat production potential and sustainability in the face of current and future global food challenges.

## Main

Bread wheat (*Triticum aestivum,* 2n=6x=42) contributes significantly to humankind’s caloric and nutritional requirements. Over the past 30 years, wheat yields have stagnated across the major growing regions suggesting that novel approaches to genetic improvement are required^1–3^. Although yield is a complex trait controlled by multiple factors, the source and sink capabilities of the plant and their interaction directly affect yield capacity^4,5^. Sink capacity involves traits influencing the size and number of sink organs, such as grains in the case of wheat, and their efficiency in mobilizing and utilizing photosynthates^5,6^. In wheat, genes controlling sink traits such as grain size have been identified and, in some cases, integrated as selection targets breeding programs ^7–15^. Another approach is to enhance sink capacity through an increase in the number of grains per wheat inflorescence, commonly referred to as the spike. This could be achieved by increasing the number of branching units along the spike, termed spikelets, which typically contain multiple florets. Alternatively, increasing the number of grains per individual spikelet is possible, albeit research is limited here ^16–19^.

A typical wheat spike contains between 20 to 28 spikelets, which each contain 3-4 fertile florets depending on the environment and genetic background^20,21^. Florets contain three stamens and a single ovary that, when fertilized, will develop into one grain^22^. There are examples of pistilloidy mutations that cause an increase in floral organ number. However, these mutations are associated with an overall reduction in fertility due to ovaries becoming stamens in some florets and stamens becoming ovaries in others, thus making these mutations not agronomically desirable^28–32^. Trigrain wheat, henceforth referred to as multi-ovary wheat, though shares some similarities with the pistilloidy mutations such as additional ovaries per floret, but importantly without the negative effect on fertility^23^. Multiovary is a strong dominant phenotype that shows high penetrance when when crossed with typical wild type wheat lines^24^. Developmental analysis of multiovary wheat line “MOV” shows significant differences in ovary development in comparison with wildtype wheat. In both genotypes, the primary ovary primordia develop first. While in wild type wheat there are no traces of any secondary ovary development, in MOV two secondary ovary primordia initiate giving rise to two ovaries which together with the primary ovary form three fully functional ovaries (Supplementary Figure 1).

### Map based cloning of *Mov-1*

Since first discovered in 1973, the underlying gene controlling the multiovary phenotype (*Mov-1*) has evaded scientists for half a century. Previous efforts mapped *Mov-1* to the distal end of chromosome arm 2DL and we reduced the interval to a 1.1 Mbp region^24–33^. In this study, we describe the use of a scaffold-assembled genome of MOV using PacBio long reads to map-based clone *Mov-1* and elucidate the gene underlying the multiovary phenotype.

For the assembly of the MOV genome we generated 212.64 Gbp of sequencing reads (∼14x coverage) and assembled them using hifiasm resulting in an assembly of 14.48 Gbp with a scaffold N50 of 15.7 Mbp. We recovered 99.4% of the Poales single copy BUSCO core genes with 96.6% coming from complete and duplicated BUSCOs (Supplementary Table 1). For precise mapping of *Mov-1,* we used an F_2_ population consisting of 1148 gametes derived from a cross between synthetic hexaploid wheat line TA8051[Prelude tetraploid/*Ae. tauschii* (TA1604)] and MOV wheat line used for the genome assembly^24^. Using four critical recombinants, we fine mapped *Mov-1* to a 134 kbp region in MOV which included two genes based on the Chinese Spring reference^34^ and *de novo* Augustus annotations (Fig. 1b,c). Interestingly, when comparing the *Mov-1* 134 kbp region with multiple wheat assemblies (e.g. Chinese Spring, 10+ Genomes, *Ae. tauschii*)^35–37^, we discovered a ∼375 kbp deletion unique to the MOV genome resulting in the deletion of ten genes (Fig. 1d; Supplementary Fig. 2).

**Figure 1.**
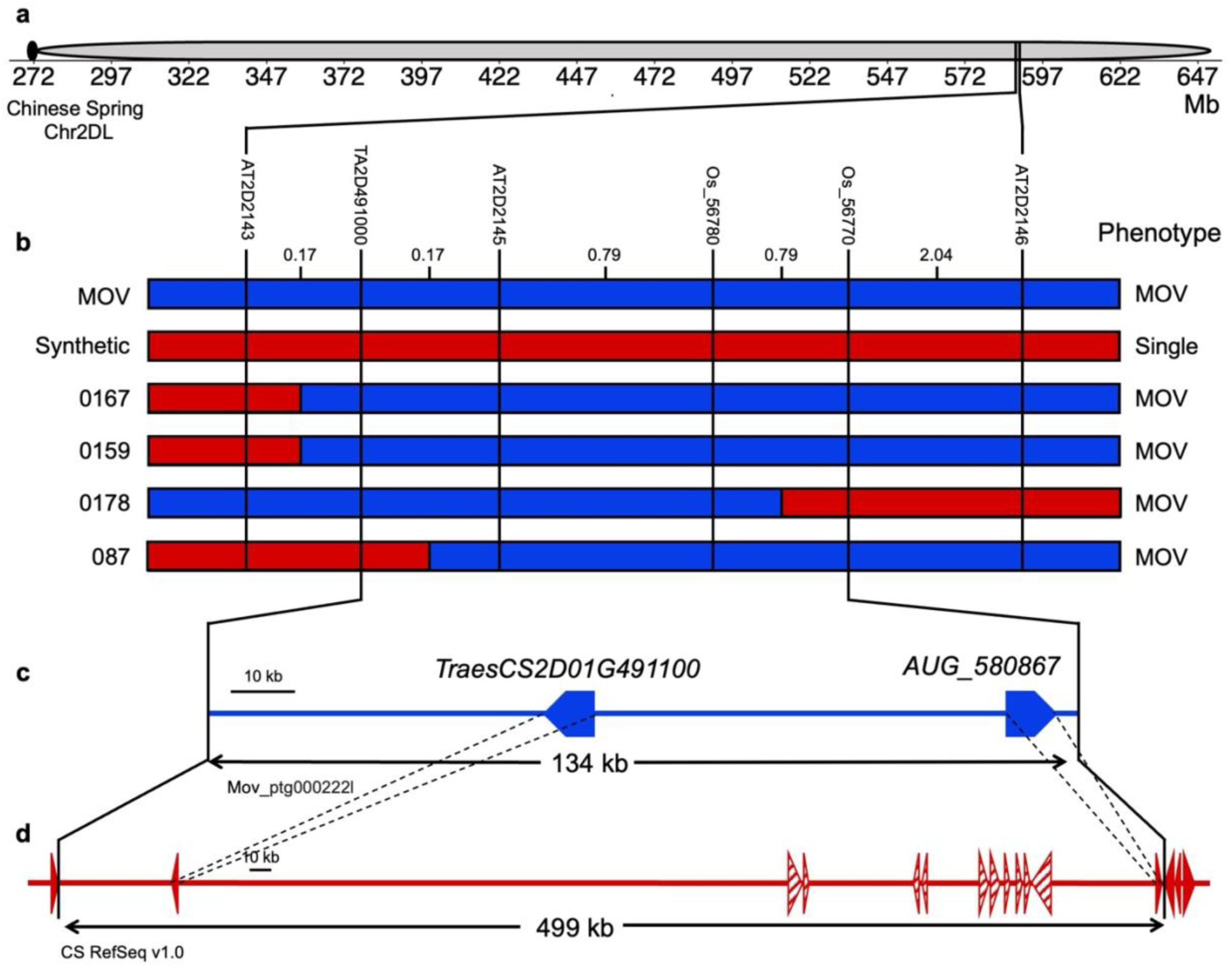
Map based cloning of *Mov-1* shows two potential candidate genes within the mapping interval. **a**. The physical location of flanking markers TA2D491000 and AT2D2146 on the long arm of chromosome arm 2DL based on RefSeq v.1.0. **b**. The genetic distance (cM) between the markers used for fine mapping as well as the recombinants used to identify the candidate region. **c**. The physical region of the fine mapping interval with the genes represented as arrows. **d.** The structure of the physical region when compared to Chinese Spring RefSeq v.1.0. Dotted lines indicate collinear genes. Dashed triangles indicate genes deleted in MOV.

To determine the putative role of these ten deleted gene on the MOV phenotype, we identified missense and truncation mutants in the Cadenza hexaploid wheat TILLING population^38^. We found mutations in seven out of the ten annotated genes within the deleted region, none of which showed any observable change in ovary number (Supplementary Table 2). For the three remaining genes that did not have mutants, we found no significant change in their expression levels based on RNA-seq data in MOV developing ovaries compared with wild type wheat line Chinese Spring. Moreover, for two deleted genes (*TraesCS2D02G491500, TraesCS2D02G492000*) we found no expression in either MOV or Chinese Spring during ovary development. This suggests that the ten deleted genes are unlikely related to the multi-ovary phenotype.

We next focused on the two genes present in the candidate region: *TraesCS2D01G491100* and *AUG_580867* ^30^. *AUG_580867*, henceforth referred to as *TaWUS2D,* encodes a homeobox containing WUSCHEL-like protein based and is orthologous to WUS-1 in *Arabidopsis thaliana* and WOX-1 in rice (*Oryza sativa),* as well as to additional WOX-family proteins in *Solanum lycopersicum, Cucumis sativus, Glycine max,* and *Citrullus lanatus*. Upregulation of WUSCHEL genes in *A. thaliana* and *C. sativus* increases floral organ number, supporting *TaWUS2* as a strong candidate gene^39–42^. We used RNAseq of developing ovaries of MOV and Chinese Spring and found that *AUG_580867* showed a 2.89-fold significantly higher expression in MOV (Table 1)^24^. To further confirm the overexpression of *TaWUS2* in MOV plants, we performed droplet digital PCR (ddPCR) using F_2:3_ lines from the MOV/TA8051 cross showing contrasting phenotypes. We found a 10.8-fold higher expression of *TaWUS2D* in developing ovaries of multi-ovary plants in comparison to single-ovary lines (Fig. 2a). We also observed significantly higher (P<0.05) expression in stem node tissue, but not in leaf tissue (Supplementary Fig 3). We performed *in-situ* hybridization in developing spikes and stem node tissues for the same lines used for ddPCR. Consistent with previous results, we found a stronger *TaWUS2D* hybridization signal in MOV across both tissues, with expression highest in developing floral tissues (Fig 2b). These results show that multi-ovary lines have higher expression of *TaWUS2D* compared to wildtype genotypes, indicating a gain of additional functionality for this putative transcription factor in the multi-ovary development.

**Figure 2.**
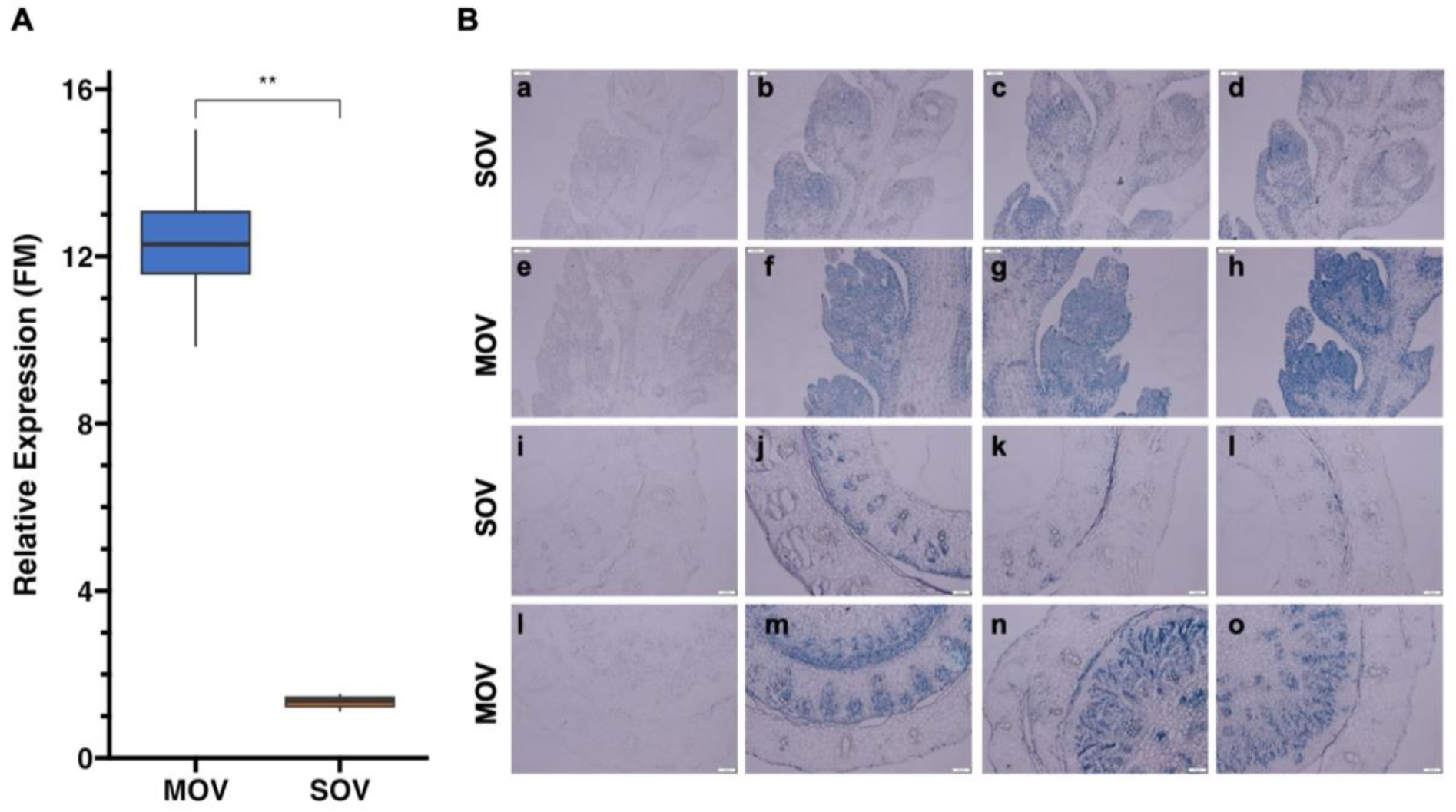
Expression patterns of *TaWUS2D.* **A**. ddPCR results of expression of *TaWUS2D* in floral meristems using the same materials as the *in situ* assay. **B. a**-**h** Developing spike tissue and **i**-**p** stem node tissue using in situ hybridization in three reps in MOV and SOV genotypes derived from a segregating F_2:3_ population between TA8051 and WT MOV, **a**, **e**, **i**, **m** are negative controls.

**Table 1.** RPKM Values and Significant Fold Changes in Genes Within the Mapping Interval.

| Gene ID | MOV | | SOV | | Fold Change<br>($p < 0.05$) |
| --- | --- | --- | --- | --- | --- |
|  | Very Young<br>Ovary (RPKM) | Young Ovary<br>(RPKM) | Very Young<br>Ovary (RPKM) | Young Ovary<br>(RPKM) |  |
| TraesCS2D02G491100.1 | 0 ± 0 | 0 ± 0 | 0 ± 0 | 0.013 ± 0.019 | ns |
| **TraesCS2D02G491200.1** | **0 ± 0** | **0 ± 0** | **0 ± 0** | **0 ± 0** | **ns** |
| **TraesCS2D02G491200.2** | **0 ± 0** | **0 ± 0** | **0 ± 0** | **0 ± 0** | **ns** |
| **TraesCS2D02G491300.1** | **0 ± 0** | **0 ± 0** | **29.315 ± 1.335** | **22.861 ± 0.076** | **-11.89** |
| **TraesCS2D02G491400.1** | **0 ± 0** | **0 ± 0** | **0.296 ± 0.098** | **0.25 ± 0.027** | **ns** |
| **TraesCS2D02G491500.1** | **0 ± 0** | **0 ± 0** | **0 ± 0** | **0 ± 0** | **ns** |
| **TraesCS2D02G491600.1** | **0 ± 0** | **0 ± 0** | **0.426 ± 0.031** | **0.378 ± 0.112** | **-8.02** |
| **TraesCS2D02G491700.1** | **0 ± 0** | **0 ± 0** | **1.835 ± 0.450** | **2.545 ± 0.003** | **-9.38** |
| **TraesCS2D02G491800.1** | **0 ± 0** | **0 ± 0** | **0 ± 0** | **0 ± 0** | **ns** |
| **TraesCS2D02G491900.1** | **0 ± 0** | **0 ± 0** | **0.025 ± 0.036** | **0.016 ± 0.023** | **ns** |
| **TraesCS2D02G492000.1** | **0 ± 0** | **0 ± 0** | **0 ± 0** | **0 ± 0** | **ns** |
| **TraesCS2D02G492100.1** | **0 ± 0** | **0 ± 0** | **0.037 ± 0.006** | **0.010 ± 0.015** | **ns** |
| *TaWUS2D(AUG_580867)* | 1.461 ± 0.268 | 0.386 ± 0.072 | 0.196 ± 0.068 | 0.122 ± 0.084 | 2.89 |
| TraesCS2D02G492200.1 | 4.385 ± 0.19 | 3.803 ± 0.155 | 3.247 ± 1.286 | 3.786 ± 0.297 | ns |

We hypothesized that sequence variation in *WUS2* cis-regulatory regions could account for the differential expression. Surprisingly, we found few and no consistent sequence variations in this region between MOV and available wildtype (single ovary) wheat assemblies (Supplementary Figure 2b). We confirmed the accuracy of the MOV assembly by sequencing a 5.7 kbp amplicon from MOV encompassing the *TaWUS2* coding region (1212 bp), 3.8 kb upstream the initiation codon and 689bp downstream the termination codon; no sequence variations were found. Additionally, we determined the 5’ and 3’ untranslated regions using rapid amplifications of cDNA ends (RACE).

### Candidate gene validation using gamma radiation induced deletions

Next, we developed a gamma radiation-induced mutant population in the MOV background to test if the deletion of *TaWUS2D* would result in loss of the MOV phenotype. Two thousand inbred MOV seeds were treated with 40Krad of gamma radiation, and surviving R₁ plants were allowed to self-pollinate. We screened the population of 358 surviving R_2_ mutants via PCR using markers spanning from 580 to 596 Mbp (encompassing *TaWUS2D* at 590.15 Mbp). We identified six independent mutants with overlapping deletions of varying size across the interval. All six mutants showed complete loss of the MOV phenotype, producing a single ovary and single seed per floret (Fig 3a-e). To determine the size of these deletions we used low-coverage whole genome sequencing of the five fertile mutants that produced seed (mutant MOV del 40-3 was lost at R_2_). The deletions ranged from ∼4 Mbp to ∼220 Mbp, consistent with the PCR screen (Fig. 3d). To further validate the loss of *TaWUS2D* we performed ddPCR for one mutant (MOV del 30-4) and *in-situ* hybridization for two mutants (MOV del 30-4, MOV del 23-1) and observed loss of expression of the *TaWUS2D* gene (Fig 3f-g). These results show that deletion of *TaWUS2D* in the MOV genetic background results in the loss of the multi-ovary phenotype.

**Figure 3.**
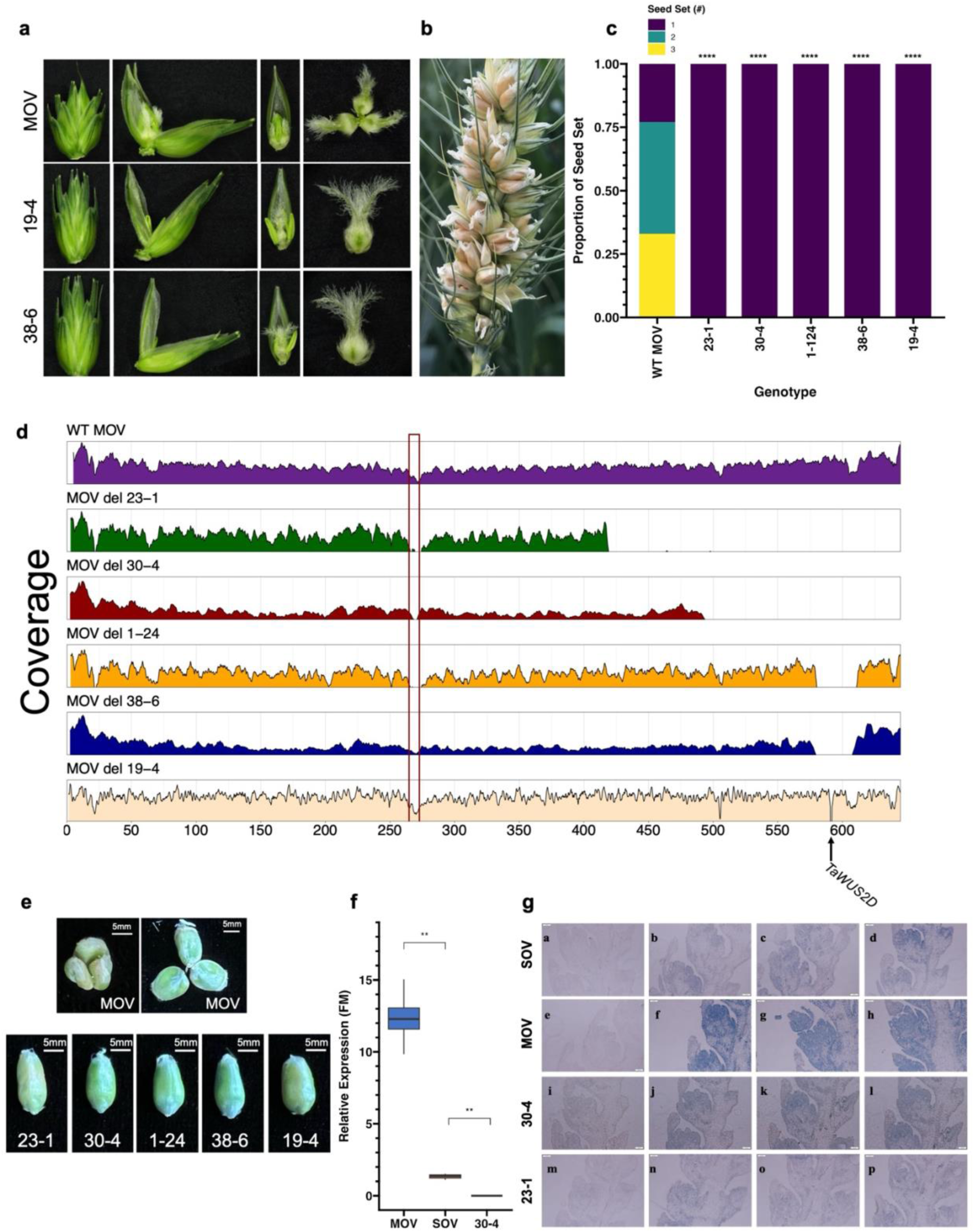
**a**. Floral phenotypes of wild type MOV and two selected deletion mutants. **b**. Close up of a MOV spike showing the grains at maturity. **c.** Proportion of number of seed set per floret in deletion mutants, **** indicates p < 0.0001 based on a non-parametric Wilcoxon test. **d.** GBS read coverage on chromosome 2D of the mutants showing deletion of the *Mov-1* genic region. The red bracket indicates the location of the centromere. **b**. **e**. Seed phenotypes in the deletion mutants. **f**. ddPCR results on MOV, SOV, and deletion mutant 30-4, ** indicate a p < 0.01 based on a student’s t-test. **g**. *In situ* hybridization of *TaWUS2D* in SOV (**a**-**d**), MOV (**e**-**f**), as well as two selected mutants 30-4 (**i**-**l**), and 23-1 (**m**-**p**), **a**, **e**, i, **m** are negative controls.

### Candidate Gene Validation Using EMS-Induced TILLING Population

Due to the unique gain-of-function mutation that gave rise to the MOV phenotype, we next set out to identify induced mutations in *TaWUS2D* using ethyl methanesulfonate (EMS) mutagenesis. We screened 1876 M_2_ mutants using markers spanning the *TaWUS2D* coding region and found 16 independent mutants, of which five mutants showed either a deleterious PROVEAN and/or SIFT score (Table 2, Fig 4a)^43,44^. Mutants with deleterious predictions were self-pollinated until the M_6_ generation to fix background mutations with the exception of mutant 555-556 which was sterile at the M_3_. The remaining mutants were phenotyped at M_6_ and M_7_ and due to the lack of a nonsense mutation, some mutants such as 555-430 and 555-641 showed some partial expression of MOV in some florets, though the presence was reduced significantly (p<0.05) in comparison with (Fig 4 b-c).

**Figure 4.**
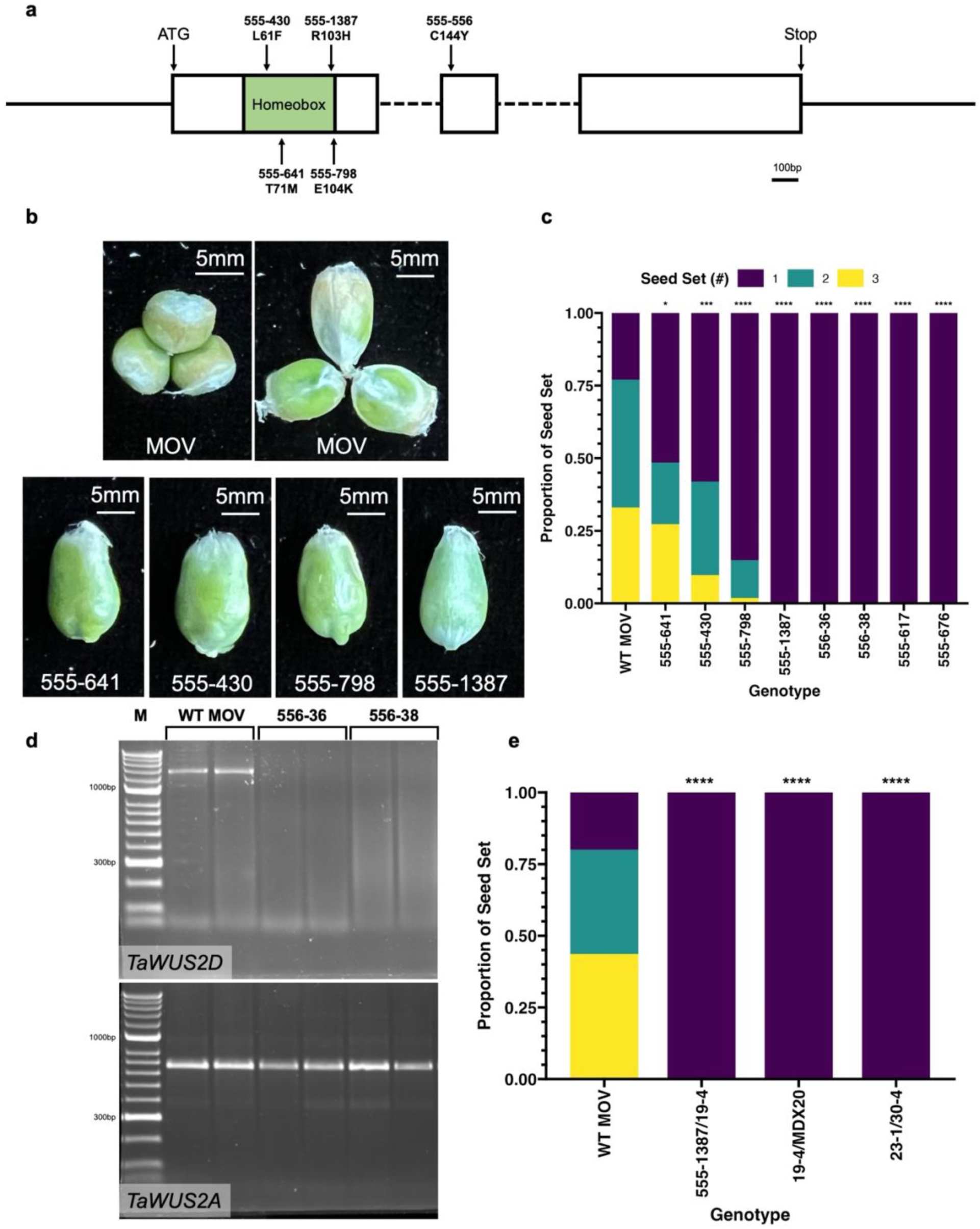
**a**. Gene structure of *TaWUS2D.* Solid lines indicate 5’ and 3’ UTRs, dotted lines indicate introns, boxes indicate exons. Arrows indicate the location of EMS mutations and amino acid changes; the red arrow indicates the mutant that was sterile and phenoytped at the M_3_ stage. **b**. Seed phenotypes during the pre-milky stage coming from EMS mutants containing point mutations. **c**. Seed set proportions in EMS mutants, * indicate p < 0.05, *** indicate p < 0.001, **** indicate p < 0.0001 based on non-parametric Wilcoxon test. **d**. Agarose gel showing lack of amplification of the *TaWUS2D* gene (Top) in two of the four EMS mutants identified in the forward screen. We used a marker designed to amplify 650 bp of the A-homeolog (*TaWUS2A*) as a control. **e**. Seed set proportions of mutant crosses, **** indicate p < 0.0001 based on non-parametric Wilcoxon test.

**Table 2.**
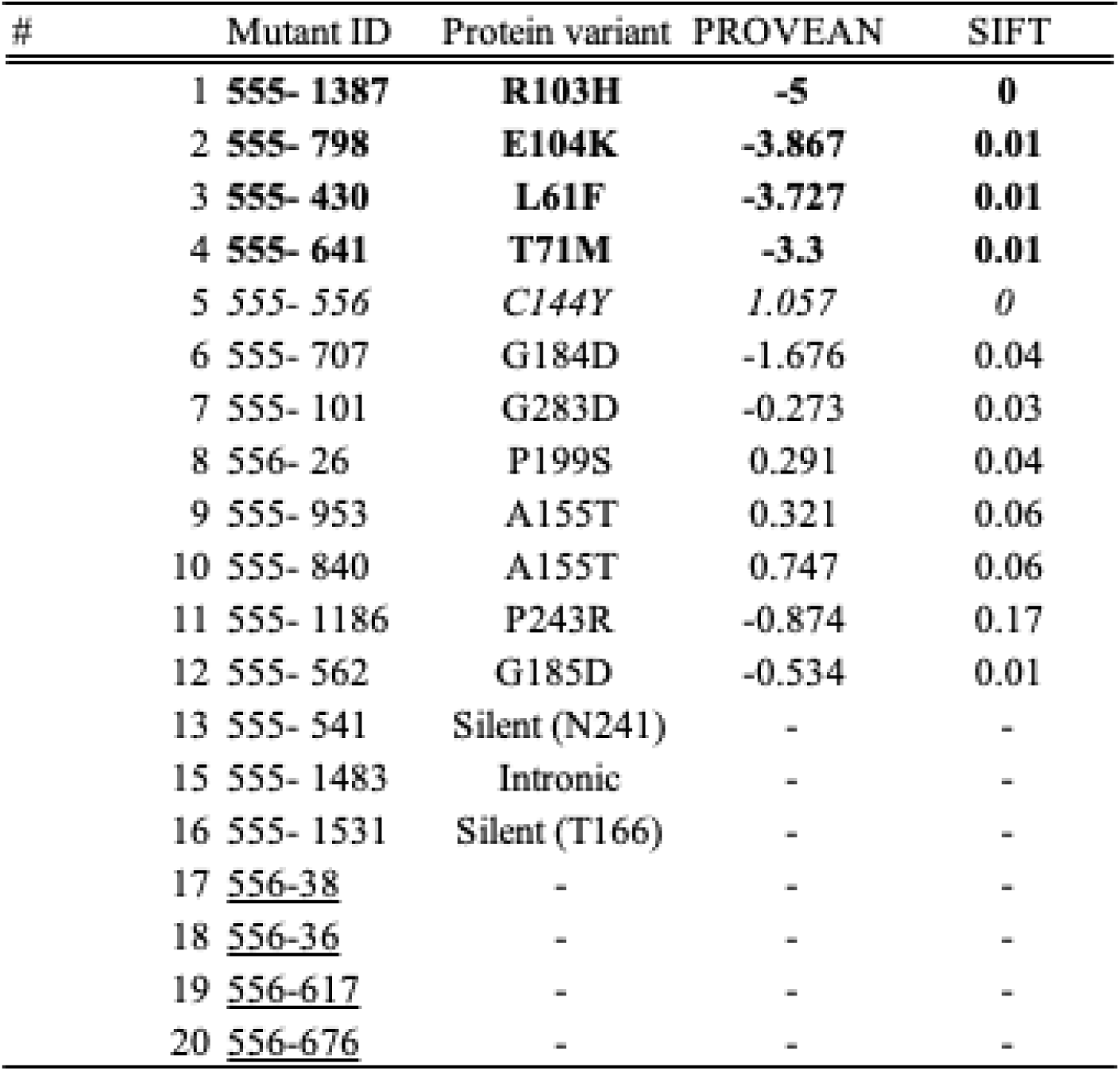
Description of TILLING Mutants Found in MOV Background. Lines in italic were lost at M_3_. Underlined lines showed complete deletion of gene.

To complement the reverse-genetics approach, we also performed a forward screen on the entire TILLING population with a total of 15,008 plants phenotyped for loss of the multi ovary trait. Along with the original mutants, we identified four additional mutants showing the single ovary phenotype. Interestingly we were unable to amplify *TaWUS2D* in these individuals, indicating a possible EMS-based deletion of this region (Fig 4d).

### Complementation of the Mutant Alleles Using Crosses of Independent Mutants

To rule out the possibility of background mutations contributing to the loss of the multi-ovary phenotype, we crossed the most significant EMS mutant (555-1387) with the deletion line with the smallest (∼4 Mbp) interstitial deletion (MOV del 19-4). Additionally, we also generated F_1_ plants by inter-crossing deletion mutants (23-1/30-4 and 19-4/38-6) and one of the deletion mutants with a single ovary bread wheat (19-4/MDX20). We observed complete loss of the multi-ovary phenotype in all crosses (Fig 4d). We also crossed deletion mutant 19-4 with wildtype MOV, and, as expected, we saw restoration of the multi-ovary phenotype in F₁ plants (Supplementary Table 3). Together, the mutant and genetic analyses show that the multi-ovary phenotype of the MOV genetic background is conditional on the presence of a functional WUS-2D gene and its protein.

## Discussion

In summary, using genetic mapping combined with reverse and forward screening of mutants in the MOV background, we identified the causal multi-ovary gene which encodes *TaWUS2D* an ortholog of WUSCHEL. This gene is upregulated by up to 10-fold in developing ovaries of multi-ovary lines compared to wild type lines. Consistent with this finding, upregulation of WUSCHEL is already known to increase carpel and locule numbers in dicot systems; here we extend its functional conservation to monocot species^12,46–49^. A question remains as to how the *TaWUS2D* over-expression phenotype evolved, as there are no sequence dissimilarities between phenotypically contrasting lines in the candidate gene protein coding or promoter regions. This suggests that nearby *cis*-regulatory sequences contribute to the ectopic upregulation of this gene. Due to the dominant gain of function of this gene in the D genome, and the seemingly developmental and tissue specific expression, simple transgenic approaches for overexpression of this gene may not be able to recreate the multi-ovary phenotype. This discovery does, however, open the door for the exploration of precise gene editing in regulatory elements to fine-tune the expression of *TaWUS2D* in single ovary wildtype lines to produce the multi-ovary phenotype. This concept of tissue specific misregulation of transcription factors based on *cis*-regulatory variation is emerging as a unifying concept for crop improvement^50–52^. The multi-ovary phenotype has excellent potential to fortify an important yield-related component by increasing the number of grains per plant, as well as has beneficial implications in hybrid-wheat by increasing threefold the number of F_1_ seed produced, thus reducing the cost per hybrid seed to compete with the cost of pure-line seeds as the cost of hybrid wheat production has become a bottleneck for their use in commercial wheat production^53–56^. It would be fascinating to see if this gene can be precisely modulated to induce specific numbers of floral organs in wheat and other monocot crop systems.

## Materials and Methods

### Plant Materials and Development of Fine Mapping Population

The MOV line of wheat was originally obtained from the Wheat Program at International Maize and Wheat Improvement Center, CIMMYT and was grown for more than five generations using single seed descent to fix any background genetic variations.

For the development of the high-resolution F_2_ population, a synthetic hexaploid wheat line TA8051 [Prelude tetraploid/Ae. tauschii (TA1604)] was used as a male pollen donor and MOV was used the female. F_1_ plants were allowed to self in order to generate a population of 1000 F_2_ plants. For fine mapping of the candidate region 574 plants (1148 gametes) were phenotyped and genotyped as described in Mahlandt et al., 2021^24^. Briefly, leaf tissue from F_2_ plants were collected and DNA was extracted. At the time of flowering, spikes were categorized as having either MOV, SOV (single ovary), or heterozygous phenotypes, depending on the developed ovaries starting from mid-spike (8-10 spikelets from the peduncle). The phenotype was confirmed at maturity by excising floret contents.

### Long-Read Sequencing and Assembly

DNA was extracted from young MOV leaf tissue roughly nine days after coleoptile emergence. Purity and concentration was confirmed using a SpectroStar Nano as well as gel electrophoresis. Library preparation and sequencing was performed using the manufacturer’s (PacBio) specifications using the PacBio Revio system.

A total of 212.64 Gb (212,637,842,631 bp) of Hifi data was generated that gave ∼14.1X coverage for the MOV genome. The data was filtered and converted to fastq using HifiAdapterFilt^57^. Hifiasm 0.18.9 was used to assemble the genome using default parameters^58^. A 14.48 GB genome was assembled with 2940 contigs. The assembled genome was scaffolded using RagTag (v2.1.0) using Triticum aestivum cv. Kariega as reference^59^. The contig level assembly was aligned to the Chinese Spring RefSeq v.1.0 reference genome using minimap2^60^. It was found that a single contig (Mov_ptg000222l, 22.7 Mb) contained our entire mapping interval. Dotplot was constructed using Chromiester between the identified contig and the Chinese Spring RefSeq v.1.0 region 587-592 Mb^61^.

### Fine Mapping of Mov-1 locus

Initial fine mapping efforts were previously completed and described by Mahlandt et al., 2021^24^. In summary, genetic markers were developed using GSP software and tested on chromosome group 2 nullisomic tetrasomic lines obtained by the Wheat Genetic Resource Center at Kansas State University (WGRC) in order to find 2D specific markers^62,63^. Markers were run on all 574 F_2_ plants. From this population, six critical recombinant lines were discovered, though they were in heterozygous condition. To confirm the interval, these six lines were advanced to F_4_ to obtain homozygous individuals (between five to nine individuals per family) and were phenotyped and genotyped again using the previously described method as described in Mahlandt et al., 2021^24^.

### Amplification and Sequencing of Mov-1

Primers were designed starting 3.8 kb upstream the start codon and 689 bp downstream the stop codon of Mov-1 and were tested for genome specificity as described prior. Using Platnum SuperFi II Master Mix (ThermoFisher 12368010) using the PCR profile suggested by the manufacturer, however 1ul of Betaine (Millipore Sigma B0300) was added to a 20ul PCR reaction. PCR products were visualized on a 1% agarose gel and bands were excised and cloned into the pCR-XL-2-TOPO vector and transformed into *E. coli* using the TOPO XL-2 Complete PCR Cloning Kit (ThermoFisher K8050-10) as per the manufacturer’s instructions.

Positive colonies were cultured in liquid media overnight and plasmid was extracted using the Zyppy Plasmid Miniprep kit (Zymo Research D4036). Plasmids were sequenced using Oxford Nanopore Technologies via Plasmidsaurus.

Additionally whole mRNA was extracted from developing MOV spiks using TRIzol reagent (Invitrogen) as per the manufacturer’s instructions and cDNA was sythesized Using AzuraQuant cDNA Synthesis Kit (Azura Genomics AZ-1995). Primers were designed flanking the coding regions of *Mov-1* and the full protein coding sequence was amplified and purified. Purified PCR products were Sanger Sequenced via GENEWIZ.

### RNA-seq Preparation and Sequencing

Ovaries were excised from MOV and an SOV wheat line Chinese Spring at very young ovary and young ovary stages and total RNA was extracted using the RNAqueous-Micro kit (ThermoFisher AM1931) and cDNAs were synthesized using the MessageAmp aRNA kit (ThermoFisher AM1751) according to the manufacturer’s instructions. Illumina RNA-seq libraries were prepared using the aRNA and the TruSeq RNA kit (version 1, rev A). Paired-end reads were obtained using the Illumina HiSeq 2000.

RNA-seq reads had adaptors trimmed and low-quality reads removed using Trimmomatic v.0.39. Clean reads were aligned to the Chinese Spring RefSeq v.1.0 using HISAT2 software using default parameters ^64^. SAM files were converted to BAM files and low-quality alignments were filtered using samtools^65^. Read counts were determined using RSubread software v. 2.16.1 using the featureCounts function and tpm and rpkm values were calculated using edgeR software v.3.42.4^66,67^. Differential expression between CS and MOV was calculated using DESeq2^68^.

### Cadenza Mutant Identification

In order to determine whether genes present the MOV native deletion play a role in the expression of the MOV phenotype we utilized the sequenced and indexed Cadenza TILLING population previously described^38,70^. Multiple deleterious mutants were identified, and their phenotypes were shared for seven of the ten genes within the region.

### 5’ and 3’ Rapid Amplification of cDNA Ends (RACE)

Rapid amplification of cDNA ends to determine the 5’ and 3’ untranslated regions (UTRs) of *Mov-1*was performed using the GeneRacer kit (Thermofisher Scientific) as per the manufacturer’s instructions. Briefly, we collected developing spike tissue from maturing MOV plants. Total RNA was isolated from this tissue using TRIzol reagent (Thermofisher scientific) following the manufacturer’s instructions. RNA quality was determine by running 1ul on a 1% agarose gel stained with ethidium bromide to ensure a lack of smearing and presence of 28S and 18S rRNA bands. Total RNA was quantified using a SPECTROstar NANO spectrophotomer (BMG LABTECH). Transcripts were amplified using a oligo-Dt primer supplied by the manufacturer and a gene-specific primer designed using the *Mov-1* sequence. Amplicons were run on a 1.5% agarose gel and bands of interest were cut and purified, and cloned using TOPO-TA cloning. Positive clones were selected, and total plasmid was purified and sequenced using Sanger sequencing. Sequences were aligned to upstream and downstream genomic regions flanking the candidate gene CDS to delineate putative UTRs.

### Development of EMS Mutant Population and Mov-1 Mutant Identification

The development of a Targeting Induced Local Lesions IN Genomes (TILLING) population and its application in hexaploid wheat is described in detail in Singh et al., 2019 ^71^. In short, to determine the LD50 for ethyl methanesulfonate treatment (EMS), 100 seeds of MOV were treated at varying concentrations EMS ranging from 0% - 1.2% and directly sown into flats. A dosage curve calculated the LD50 at 0.6%, and ∼4000 seeds (by weight) of MOV were treated at this concentration and directly sown into plots in greenhouse settings. The M_1_ population was allowed to self to give rise to an M_2_ population of 1876 individuals. At the two-leaf stage, leaf tissue was harvested from each individual plant and high-throughput DNA extraction was performed. DNA samples were normalized to 20ng/ul for downstream analysis. A subset of DNA for each sample were bulked to create 4X pools, retaining the sample’s plate position.

Several markers were developed to span the length of the *Mov-1* candidate gene in order to discover deleterious SNPs within the coding region and tested for specificity as described previously. Markers were run on the 4X pools first and SNPs were determined using the Cel-1 assay as described in Singh et al., 2019. Positive mutant pools were then deconvoluted to identify individuals harboring SNPs. PCR products from mutants were purified and Sanger sequenced using the BigDye Terminator v3.1 Cycle Sequencing Kit (ThermoFisher, cat. 4337455) and the 3730xl DNA Analyzer (ThermoFisher, cat. A41046) as per the manufacturer’s instructions. Sequences obtained from the DNA analyzer were aligned to the wild-type sequence using SnapGene v6.1. Mutation effects were calculated using Protein Variation Effect Analyzer (PROVEAN) v1.1 software as well as Sorting Intolerant from Tolerant (SIFT) software^43,44^.

A total of four mutants with both deleterious SIFT and PROVEAN scores were identified within the population and were allowed to self until M_6_ generation in order to fix any background mutations. For two subsequent generations (M_6_ and M_7_) 15 plants were phenotyped per mutant at maturity for ovary number and seed set. Phenotypes were collected starting at the center spikelet with one lateral floret chosen and moving one spikelet up and one spikelet down on either side resulting in six florets phenotyped per plant. Seed phenotypes were considered as either one, two, or three seeded.

### Development of MOV Deletion Panel and Genotyping

In order to create a deletion panel in the background of MOV, 2000 seeds from inbred MOV were plated on petri dishes and sent to the Oregon State University Radiation Center. Plates were spaced evenly in a Gammacell 220 60Co gamma irradiator and treated with 40Krad of gamma radiation. After treatment, R1 seeds were directly sown into pots. Due to the high radiation treatment, only 358 R2 plants were able to be obtained. Tissue from R2 plants were collected at the two-leaf stage and DNA was extracted.

Molecular markers were designed and tested for specificity spanning from chromosome 2D:580Mb-596Mb with 2Mb intervals between each marker. The deletion panel was genotyped using these markers as well as a gene marker for *Mov-1*. Seven independent lines coming from separate families showed deletions within the *Mov-1* region as well as the flanking marker, and these plants were advanced to R_5_ through single seed descent to fix any background deletions.

To understand the sizes of these deletion on a chromosome scale we chose to perform short-read whole-genome sequencing (WGS) using genotyping by sequencing (GBS)^72^. Using a modified method described by Singh et al., 2020, DNA was digested by Pst1 and Msp1 restriction endonucleases and barcode adaptors were ligated to the digested products. Libraries constructed and sequenced using the NOVASEQ 6000 system at 384-plex. Reads obtained were demultiplexed as well as low-quality filtered and adaptor trimmed using Ultraplex software v.1.2.10^73^. The trimmed reads were then aligned to the Chinese Spring reference v.1.0 genome using Bowtie2 v.2.5.3 software using the following parameters: --threads 40 -q --end-to-end -D 20 -R 3 -N 0 -L 10 -i S,1,0.25. SAM files were converted to BAM files using samtools as mentioned previously. Coverage was calculated for chromosome 2D using Bedtools software v.2.30.0 with the command bedtools coverage at a sliding window of 500kb ^74^. Coverage was then visualized using R software v.4.3.1. Similar to the EMS mutants, R_5_ and R_6_ generations of plants were phenotyped in the same method as described previously.

### Development of F_1_ Mutant Crosses

To further delineate the physical location of the *Mov-1* gene, multiple crosses were made between mutants. First a cross was made between MOV deletion mutant 19-4 and a SOV line soft red winter wheat (SRWW) Maryland variety MDX20. In this cross mutant 19-4 was used as the female and MDX20 as the pollen source. Next, a cross was made between two of the MOV deletion lines harboring the largest deletions (MOV del 30-4 and MOV del 23-1) with MOV del 23-1 used as the female and MOV del 30-4 used as a pollen source. Finally, to localize the mutation to the *Mov-1* gene, a cross was made between the EMS mutant with the lowest PROVEAN and SIFT scores as well as the strongest loss of phenotype (555-1387) and the MOV deletion line with the smallest (∼4 Mb) deletion (MOV del 19-4), using 555-1387 as the female and MOV del 19-4 as the pollen source.

F_1_ seeds for the MDW20/19-4 were sown first and given 4 weeks of vernalization as it is a cross between a spring wheat line and a winter wheat line. Ten days before MDW20/19-4 plants came out of vernalization, the remaining crosses as well as WT MOV were sown. After vernalization, all plants were moved to a growth chamber and allowed to grow to maturity. Phenotyping of these plants were done in the same manner as previously described.

### In-situ Hybridization to Assay Localized Gene Expression

In situ PCR was performed following the protocol described previously with modifications [62]. Spikes and stem nodes at the spikelet differentiation stage were sampled from an SOV and MOV lines derived from a F2:3 line of MOV/Synthetic showing single-ovary and multi-ovary phenotypes respectively, and MOV deletion lines 23-1 and 30-4 and fixed with fresh-prepared PFA solution (2.5% glutaraldehyde and 4% paraformaldehyde in 1× PBS) at 4°C overnight. The fixed tissues were washed with 1×PBS and dehydrated using a graded ethanol series. Subsequently, the tissues were embedded in paraffin. The spike tissues were sectioned longitudinally and stem nodes transfersely at a thickness of 10μm using a histology microtome (Leica Biosystems) and mounted on precleaned glass slides placed on a 45°C hotplate. Glass slides containing tissue 140 sections were deparaffinized with xylene, rehydrated through a graded ethanol series, and then air-dried. After post fixation with 4% PFA for 4 h at room temperature, slides were washed twice with 1XPBS for 10 min and digested with proteinase K for 5–60 min at 55°C. Subsequent treatments, including DNase treatment, in situ reverse transcription, in situ PCR, and colorimetric detection of digoxin (DIG)-labeled PCR products, were performed on slides placed in Frame-Seal incubation chambers. DNase treatment was proceeded with TURBOTM DNase (Invitrogen).

In situ reverse transcription was performed on the DNase-treated slides using the Affinity Script QPCR cDNA Synthesis Kit (Agilent Technologies) based on the manufacturer’s instructions. For in situ PCR reaction, 200 μL PCR reaction solution containing gene-specific primers, Phusion High-Fidelity DNA Polymerase (Invitrogen catalog number F530S) and 4 μM DIG-11-dUTP (Sigma-Aldrich catalog number 11093088910) were applied on the slides with tissues, slides were sealed with a frame-seal chamber and incubate in a thermocycler with an optimized program. Colorimetric detection of DIG-labeled PCR products was performed with Anti-DIG-AP (Sigma-Aldrich catalog number 11093274910), followed by staining with BM-purple (Sigma-Aldrich catalog number 11442074001). The sections were visualized under a bright field microscope and images were taken using a high-resolution K8 camera attached to the Leica Thunder microscope. Negative controls were performed and analyzed using sections from the same tissue samples processed as described above, with the exception that the in situ reverse transcription step was omitted.

### Droplet Digital PCR Assay

To quantify the expression of mRNAs this study, the droplet digital PCR (ddPCR) assay was applied. The primers and probes were designed correspondingly to target different genes or miRNAs. The SNPs and indels between wheat homeologs were considered to increase the specificity of the primers and probes for A,B, D genomic copies, and an intron-spanning feature was included in the primers to eliminate off-target binding to potential genomic DNA contaminations for detection and measuring targeted gene expression levels and profiles. The primers and probes for targeting wheat ACTIN2 were also designed as the reference controls for the expression of genes.

Total RNA was isolated using TRIzol reagent (Invitrogen) with the tissues of wheat collected as described above. Two micrograms of total RNA isolated was treated with DNase I (ThermoFisher 18068015), then used as template for reverse transcription (RT) with the Affinity Script QPCR cDNA Synthesis Kit (Agilent 600559) based on the manufacturer’s instructions. For ddPCR assay, The probes were labeled at the 5’ end with either 6-carboxyfluorescein or 6-carboxy-2,4,4,5,7,7-hexachlorofluorescein succinimidyl ester as the reporter, and labeled with ZEN and Iowa Black FQ at the 3’ end as the double quenchers (Integrated DNA Technologies). For gene expression analyzed by Droplet Digital PCR, 20 μL volume reaction system containing ddPCR SuperMix for probes (no dUTPs, Bio-Rad 1863024), cDNA templates, forward and reverse primers and specific probes with optimized concentration were mixed with 70 μL of Droplet Generation oil for Probes in a DG8 Cartridge (Bio-Rad 1864008) loaded into the QX200 Droplet Generator (Bio-Rad) to generate PCR droplets. By the end of the droplet generation, 40 μL of each droplet mixture was transferred to a 96-well PCR plate and sealed with a PX1TM PCR Plate Sealer (Bio-Rad). PCR thermal cycling was optimized, and amplification signals were read using the QX200TM Droplet Reader and analyzed using QuantaSoft software (Bio-Rad). Three biological replicates were performed for each experiment.

