## Supplementary Data for "*TaWUS2D* regulates the number of grains per spikelet by enhancing the number of fertile ovaries in Multi-Ovary Wheat"

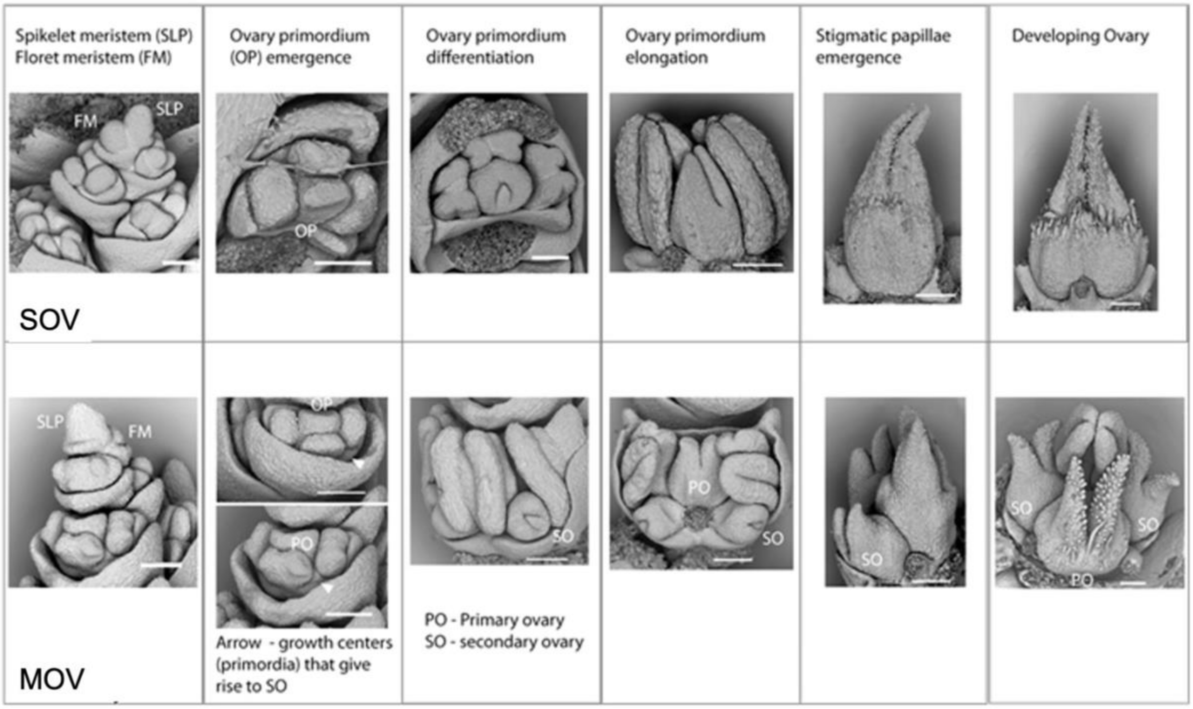


Supplementary Figure 1. SEM figures showing developing floral organs in SOV (top) and MOV (bottom).

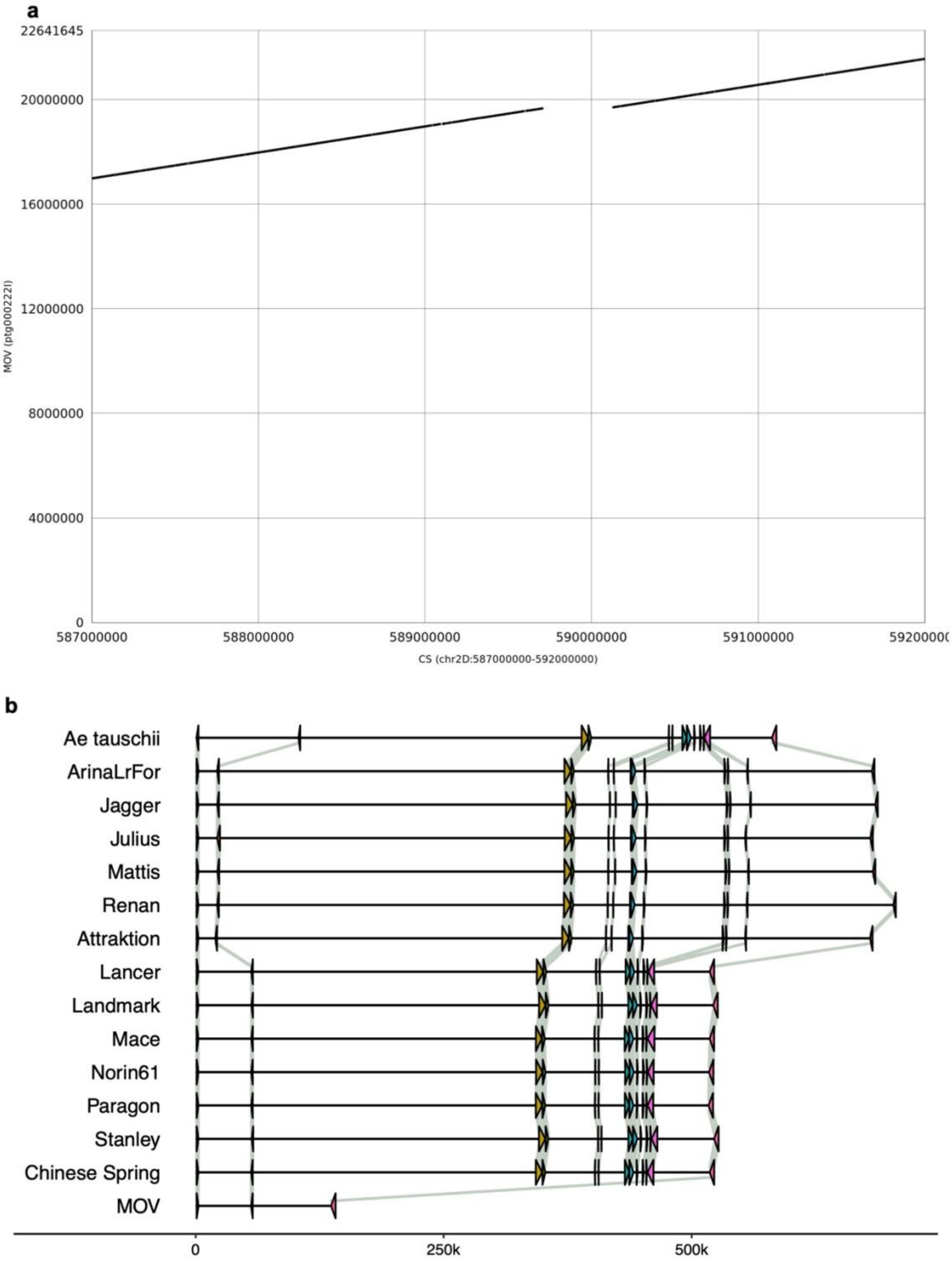

a
22641645
20000000
16000000
12000000
8000000
4000000
0
MOV (ptg000222l)
587000000
588000000
589000000
590000000
591000000
CS (chr2D:587000000-592000000)
b
Ae tauschii
ArinaLrFor
Jagger
Julius
Mattis
Renan
Attraktion
Lancer
Landmark
Mace
Norin61
Paragon
Stanley
Chinese Spring
MOV
0
250k
500k

Supplementary Figure 2. Exploration of the genomic architecture around the Mov-1 locus. **a.** A dotplot showing collinearity of the scaffold used for fine mapping *Mov-1* (Mov_ptg000222l) and the corresponding coordinates of Chr2D of Chinese Spring RefSeq v1.0. showing a large deletion event. **b.** Comparison of the fine-mapping interval in MOV background compared to the 10+ genomes as well as *Ae. tauschii* showing that this event is unique to this genotype.

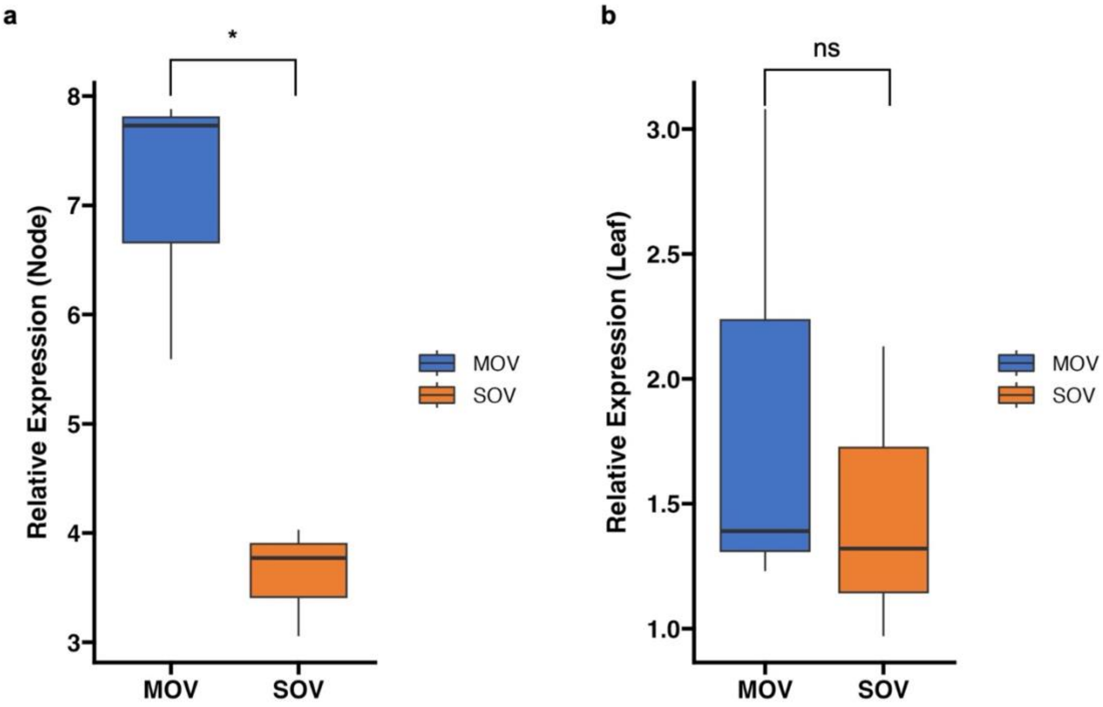


Supplementary Figure 3. **a.** ddPCR results of leaf tissue using the same plant material described earlier showing no significant change in expression between MOV and SOV. **b.** ddPCR results of node tissue showing a significant ($p<0.05$) upregulation in MOV when compared to SOV.

Supplementary Table 3. Mutant/Deletion Cross Information Describing Cross Type, Expected, and Observed Phenotypes Isolating Mutant Allele Effect on MOV Phenotype.

Supplementary Table 1. Statistics of the Long-Read Assembly of MOV Wheat

| Parameter | Value |
|---|---|
| **Total Contigs** | 2940 |
| **Genome Size (Mb)** | 14479.711 |
| **Contig N/L50 (Mb)** | 241/15.652 |
| **Max scaffold length (Mb)** | 117.889 |
| **Complete BUSCOs (%)** | 99.39 |
| **Complete and single-copy BUSCOs (%)** | 2.82 |
| **Complete and duplicated BUSCOs (%)** | 96.57 |
| **Fragmented BUSCOs (%)** | 0.06 |
| **Missing BUSCOs (%)** | 0.55 |
| **Total BUSCO groups searched** | 4896 |

Supplementary Table 2. Mutants Identified and Their Phenotypes in cv. Cadenza Background for Genes Present in MOV Native Deletion

| Gene ID | Mutant ID | Mutantion Type | AA Change | Phenotype |
|---|---|---|---|---|
| TraesCS2D02G491200 | Cadenza0161 | missense variant | P/S | **Flag leaf senescence before 24.6** |
| | Cadenza0911 | missense variant | G/D | No observable Phenotype |
| | Cadenza0240 | missense variant | G/E | No observable Phenotype |
| | Cadenza1273 | missense variant | P/S | No observable Phenotype |
| | Cadenza0469 | missense variant | R/Q | No observable Phenotype |
| | Cadenza0604 | missense variant | H/Y | No observable Phenotype |
| | Cadenza0474 | missense variant | P/S | No observable Phenotype |
| | Cadenza0630 | missense variant | G/E | No observable Phenotype |
| | Cadenza1715 | missense variant | T/I | No observable Phenotype |
| | Cadenza1677 | missense variant | A/V | No observable Phenotype |
| | Cadenza1070 | missense variant | P/L | **Small** |
| | Cadenza1599 | missense variant | R/C | **extra spikelet, reduced, #8** |
| | Cadenza0348 | missense variant | P/L | No observable Phenotype |
| | Cadenza1517 | missense variant | E/K | No observable Phenotype |
| | Cadenza1270 | missense variant | L/S | **Fat ears** |
| | Cadenza1443 | missense variant | C/Y | No observable Phenotype |
| | Cadenza0161 | missense variant | P/S | **Flag leaf senescence before 24.6** |
| | Cadenza0911 | missense variant | G/D | No observable Phenotype |
| | Cadenza0240 | missense variant | G/E | No observable Phenotype |
| | Cadenza1273 | missense variant | P/S | No observable Phenotype |
| | Cadenza0469 | missense variant | R/Q | No observable Phenotype |
| | Cadenza0604 | missense variant | H/Y | No observable Phenotype |
| | Cadenza0474 | missense variant | P/S | No observable Phenotype |
| | Cadenza0630 | missense variant | G/E | No observable Phenotype |
| | Cadenza1715 | missense variant | T/I | No observable Phenotype |
| | Cadenza1677 | missense variant | A/V | No observable Phenotype |
| | Cadenza1070 | missense variant | P/L | **Small** |
| | Cadenza1599 | missense variant | R/C | **extra spikelet, reduced, #8** |
| | Cadenza0348 | missense variant | P/L | No observable Phenotype |
| | Cadenza1517 | missense variant | E/K | No observable Phenotype |
| | Cadenza1270 | missense variant | L/S | **Fat ears** |
| | Cadenza1443 | missense variant | C/Y | No observable Phenotype |
| TraesCS2D02G491300.1 | Cadenza0155 | missense variant | P/L | No observable Phenotype |
| | Cadenza0553 | missense variant | A/V | **Stripes/spots** |
| | Cadenza1392 | missense variant | G/E | No observable Phenotype |
| | Cadenza0899 | missense variant | T/I | **Stripes/spots** |
| TraesCS2D02G491600.1 | Cadenza0477 | missense variant | G/E | No observable Phenotype |
| | Cadenza0562 | missense variant | N/S | No observable Phenotype |
| | Cadenza0019 | missense variant | N/S | No observable Phenotype |
| | Cadenza2098 | missense variant | N/S | No observable Phenotype |
| | Cadenza1044 | missense variant | N/S | No observable Phenotype |
| | Cadenza1567 | missense variant | N/S | No observable Phenotype |
| | Cadenza1605 | missense variant | N/S | No observable Phenotype |
| | Cadenza0159 | missense variant | N/S | No observable Phenotype |
| | Cadenza2070 | missense variant | N/S | No observable Phenotype |
| | Cadenza2107 | missense variant | N/S | No observable Phenotype |
| | Cadenza0877 | missense variant | N/S | No observable Phenotype |
| | Cadenza0887 | missense variant | N/S | No observable Phenotype |
| | Cadenza0392 | missense variant | N/S | No observable Phenotype |
| | Cadenza1714 | missense variant | N/S | No observable Phenotype |
| TraesCS2D02G491700.1 | Cadenza1746 | missense variant | P/L | No observable Phenotype |
| | Cadenza1478 | missense variant | G/V | **Flag leaf senescence before 1.7** |
| | Cadenza0156 | missense variant | G/E | No observable Phenotype |
| | Cadenza0871 | stop gained | Q/* | No observable Phenotype |
| | Cadenza1069 | missense variant | R/Q | No observable Phenotype |
| TraesCS2D02G491800.1 | Cadenza1535 | stop gained | L/* | No observable Phenotype |
| | Cadenza1375 | stop gained | L/* | **Flag leaf senescence before 1.7 Few** |
| | Cadenza1793 | stop gained | L/* | No observable Phenotype |
| | Cadenza0999 | stop gained | L/* | No observable Phenotype |
| | Cadenza1715 | stop gained | L/* | No observable Phenotype |
| | Cadenza0167 | stop gained | L/* | **BIG** |
| | Cadenza1987 | stop gained | L/* | No observable Phenotype |
| | Cadenza0450 | missense variant | A/T | No observable Phenotype |
| TraesCS2D02G491900.1 | Cadenza1800 | missense variant | A/T | No observable Phenotype |
| | Cadenza0393 | missense variant | S/P | **Flag leaf senescence before 24.6** |
| | Cadenza0620 | missense variant | G/W | No observable Phenotype |
| | Cadenza1448 | missense variant | G/W | No observable Phenotype |
| | Cadenza1762 | missense variant | A/T | No observable Phenotype |
| | Cadenza1713 | missense variant | C/Y | No observable Phenotype |
| TraesCS2D02G492100.1 | Cadenza1816 | missense variant | A/V | No observable Phenotype |
| | Cadenza1052 | missense variant | R/C | No observable Phenotype |
| | Cadenza0230 | missense variant | P/S | No observable Phenotype |
| | Cadenza1750 | missense variant | P/S | **Stripes/spots** |
| | Cadenza0669 | missense variant | E/K | No observable Phenotype |
| | Cadenza1773 | missense variant | A/T | No observable Phenotype |

Supplementary Table 3. Mutant/Deletion Cross Information Describing Cross Type, Expected, and Observed Phenotypes Isolating Mutant Allele Effect on MOV Phenotype.

| # | Cross | Generation | Cross Type | Expected Phenotype | Observed Phenotype |
|---|---|---|---|---|---|
| 1 | 555-1387/19-4 | $F_1$ | EMS/Deletion | SOV | SOV |
| 2 | 19-4/MDX20 | $F_1$ | Deletion/SOV | SOV | SOV |
| 3 | 23-1/30-4 | $F_1$ | Deletion/Deletion | SOV | SOV |
| 4 | 19-4/38-6 | $F_1$ | Deletion/Deletion | SOV | SOV |
| 5 | 19-4/MOV WT | $F_1$ | Deletion/MOV | MOV | MOV |
